# Vampire bats in Belize harbor multiple *Trypanosoma cruzi* genotypes: implications for parasite transmission at the wildlife–domestic–human interface

**DOI:** 10.64898/2025.12.10.693612

**Authors:** Annalise Dunsmore, Kristin E. Dyer, Lauren R. Lock, Weihong Tu, M. Brock Fenton, Nancy B. Simmons, Eric Dumonteil, Daniel J. Becker, Claudia Herrera

## Abstract

**Background:** Chagas disease, caused by *Trypanosoma cruzi*, is a neglected tropical disease with complex sylvatic and domestic transmission cycles involving vectors, mammalian hosts, and humans. The common vampire bat (*Desmodus rotundus*) is an obligate blood-feeding species that frequently interacts with livestock and humans, yet their role in parasite maintenance in Central America remains poorly characterized.

**Methodology/Principal Findings:** We analyzed 205 blood samples from vampire bats collected at two sites in northern Belize over three years (2019, 2021, 2022). PCR screening revealed an overall *T. cruzi* prevalence of 41.5% and increasing infection risks over time. The amplicon-based next-generation sequencing of the parasite mini-exon locus identified 36 unique haplotypes belonging to five DTUs: TcI, TcIV, TcV, TcVI, and TcBat. TcBat was present in half of the samples sequenced.

TcBat and TcVI, were detected in Belize for the first time. Belizean TcBat haplotypes clustered closely with Colombian reference sequences, and a subset formed a Belize-specific clade, indicating previously unrecognized TcBat diversity in the region; in contrast, Brazilian TcBat sequences were more distantly related.

**Conclusions/Significance:** Vampire bats in Belize harbor diverse *T. cruzi* genotypes, including DTUs with established roles in human disease. Given their obligate blood diet, frequent feeding on livestock, and occasional biting of humans, vampire bats may serve as bridge hosts linking sylvatic, domestic, and human transmission cycles. Together with a recent report of an acute human Chagas disease case in northern Belize, these results underscore the need for integrated One Health surveillance of bats, vectors, livestock, and humans to better evaluate and mitigate the risk of Chagas disease in Central America.

**Author Summary:** Chagas disease, caused by the parasite *Trypanosoma cruzi*, is a major public health concern in the Americas but remains neglected in many regions, including Central America. The parasite is typically transmitted by kissing bugs, but wild mammals also serve as important hosts. Vampire bats (*Desmodus rotundus*) are unique because they feed exclusively blood and frequently bite livestock, and occasionally humans, creating opportunities for parasite transmission across different environments. In this study, we screened 205 vampire bats from northern Belize and found that over 40% were infected with *T. cruzi*. By sequencing parasite DNA, we discovered a surprising diversity of DTUs, including TcI, TcIV, TcV, TcVI, and TcBat. The detection of TcVI is particularly important, because this DTU, associated with human infections in South America, had not been previously reported in Belize. TcBat haplotypes were highly prevalent and genetically diverse. Our results show that vampire bats are important hosts of diverse *T. cruzi* genotypes in Belize and may act as bridge hosts between wildlife, livestock, human transmission cycles. Enhanced One Health surveillance across vectors, bats, domestic animals, and humans will be critical for understanding and preventing Chagas disease emergence in Central America.

## Introduction

Chagas disease, caused by the protozoan parasite *Trypanosoma cruzi*, is a major neglected tropical disease in the Americas, affecting an estimated 6–7 million people and placing nearly 70 million at risk of infection [1, 2]. Clinically, infection can lead to chronic cardiac, gastrointestinal, and neurological complications, which together make Chagas disease a leading cause of cardiomyopathy in Latin America [3]. The parasite occurs from the southern United States to Argentina and Chile, reflecting its remarkable ecological adaptability and capacity to persist across diverse host and vector species [1, 4]. Transmission occurs through multiple routes, including vector-borne, oral, congenital, and transfusional pathways [4] that interconnect mammalian hosts and insect vectors, creating highly dynamic eco-epidemiological networks [5].

*T. cruzi* is genetically diverse, comprising seven discrete typing units (DTUs: TcI–TcVI and TcBat) that differ in their geographic distribution, host associations, and clinical relevance [6, 7]. This remarkable heterogeneity is shaped by selective pressures from a wide range of mammalian hosts and vectors, resulting in profound biological and epidemiological differences among DTUs [8]. Bats are particularly important in this context: phylogenetic studies support the “bat-seeding” hypothesis, whereby the common ancestor of *T. cruzi* emerged from a bat trypanosome, with TcI representing the first major lineage to expand into terrestrial mammals[9, 10]. While studies in Central America have rarely documented *T. cruzi* infection in bats [11], evidence from Colombia and South America supports the role of these flying mammals in maintaining an extensive genetic diversity[12–14]. In Colombia, DTUs such as TcI and TcBat have been detected in frugivorous and insectivorous bats, suggesting active transmission cycles involving diverse chiropteran hosts [12]. Similarly, research in Ecuador [13] and Brazil has shown that bats harbor multiple *Trypanosoma* species and contribute to the maintenance and dispersal of *T. cruzi* in fragmented landscapes [14]. The common vampire bat (*Desmodus rotundus*) is of particular concern because of its obligate hematophagy and frequent feeding on both livestock and humans [15, 16], positioning it as a potential bridge host linking sylvatic and domestic cycles of parasite transmission.

Despite the ecological importance of bats, their role in *T. cruzi* transmission in Central America remains especially poorly understood. In Belize, *Triatoma dimidiata* sensu lato has historically been considered the primary vector of *T. cruzi*, with most populations found in sylvatic habitats and only seasonal household infestations reported in the Cayo and Toledo districts in the central and southern region of the country [17]. However, recent findings challenge the assumption of limited transmission risk in the north of Belize. In 2020, the first confirmed autochthonous case of acute Chagas disease in Belize was diagnosed in a child from Sarteneja, in the Corozal District. Molecular analysis revealed a multiclonal infection involving TcII, TcIV, and TcV DTUs, and triatomine vectors were found within the household, one of which tested positive for *T. cruzi* [18]. Subsequent entomological investigations identified a novel *Triatoma* species closely related to *T. huehuetenanguensis* in the peridomestic environment of the same household. This vector also harbored *T. cruzi* TcIV, genetically matching the strain found in the child, suggesting active local transmission and the presence of previously unrecognized vector diversity [19].These findings underscore the need to reassess the transmission ecology and diversity of *T. cruzi* in northern Belize.

Here, we used molecular diagnostic and amplicon-based next-generation sequencing (NGS) to investigate *T. cruzi* infection prevalence and genetic diversity in *D. rotundus* populations in the Orange Walk District of northern Belize. By characterizing parasite genotypes at high resolution, our goal was to elucidate the role of vampire bats in local transmission cycles and assess the implications for public health in this understudied region of Central America.

## Methods

### Study area and bat sampling

As part of a long-term study on vampire bat ecology and pathogen dynamics [20–23] , we analyzed blood samples from *Desmodus rotundus* collected during three field seasons: April 2019 (n = 136), November 2021 (n = 14), and April 2022 (n = 55). Sampling was conducted at two sites in the Orange Walk District of northern Belize, the Lamanai Archaeological Reserve (LAR) and Ka’Kabish (KK), which are separated by approximately 8 km and embedded in a heterogeneous landscape of agriculture, pasture, and forest fragments [24, 25]. The LAR represents a minimally disturbed 450-ha semi-deciduous tropical forest, while KK is a 45-ha fragment of secondary growth forest (Figure 1).

**Figure 1.**
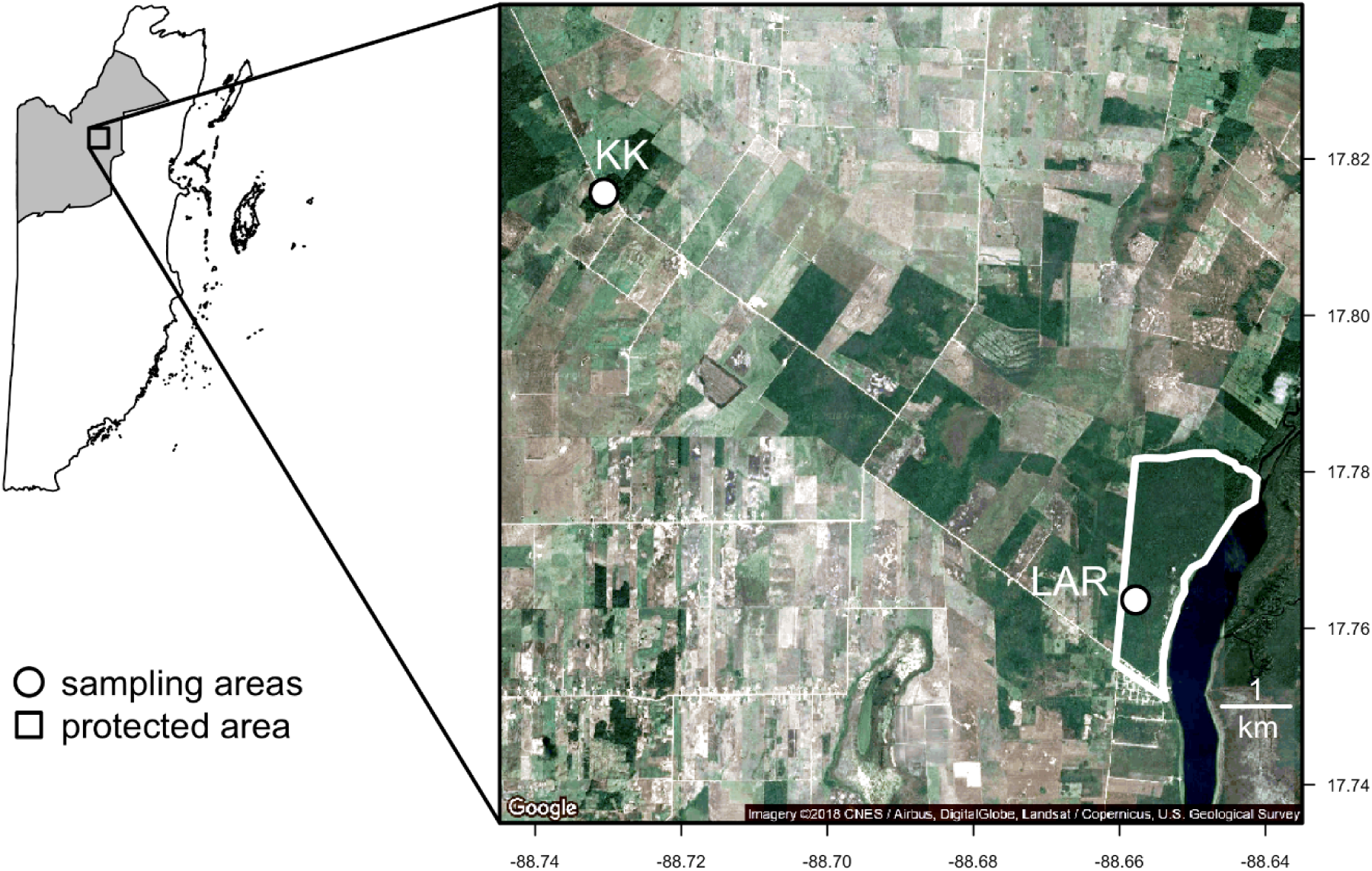
Map of the two study sites in northern Belize where samples were obtained. Borders (in color) mark the boundaries of Ka’Kabish (KK) and the Lamanai Archaeological Reserve (LAR).

Bat captures were conducted during the dry seasons (2019, 2022) and the rainy season (2021). Roost abandonment prevented sampling at KK in 2021. We captured bats using harp traps and mist nets positioned at roost exits and along flyways. Individuals were marked with 3.5 mm incoloy wing bands (Porzana Inc.), and sex, age class, and reproductive status were recorded. Blood was obtained by lancing the propatagial vein with a 23-gauge needle, followed by collection in heparinized capillary tubes. Blood was stored on Whatman FTA cards with desiccant, stored at ambient temperature in the field, and later kept at −20 °C at the University of Oklahoma.

Bleeding was stopped using styptic gel, and bats were released at the capture site. All animal handling protocols followed guidelines of the American Society of Mammalogists and were approved by the AMNH (IACUC 20190129) and the University of Oklahoma (R21-021). Bat captures were authorized by the Belize Forest Department (FD/WL/1/19(09), FD/WL/1/21(12)) and the Belize Institute of Archaeology (IA/S/5/6/21(01)).

### Trypanosoma cruzi *diagnostics*

We extracted genomic DNA from blood on FTA cards using QIAamp DNA Investigator Kits (Qiagen) as described previously, with minor modifications to the manufacturer protocols [26, 27]. Two standard detection PCR assays using previously described primers, targeting a 330 bp minicircle kinetoplast DNA sequence (121/122) and a 188 bp fragment of a highly repetitive genomic satellite DNA present in *T. cruzi* (TCZ1/TCZ2), respectively [28, 29], were performed on all DNA samples. PCR products were then subjected to gel electrophoresis on an ethidium bromide–stained 2.0% agarose gel run for 50 minutes at 100 V to determine infection status. Samples were considered diagnostically positive for parasite presence if at least one of the two PCR reactions showed a positive result indicated by presence of a 188 bp or 330 bp band for the respective reactions. Extraction controls (blank punches from FTA cards) and negative controls (molecular-grade water) were included in all PCR reactions. In addition, positive control from culture, *T. cruzi* strain SC43, a well-characterized strain representative of a TcV DTU, at 10 fg/µL was included in diagnostic PCR reactions [30]. A small subset of 2019 *T. cruzi* positivity data (*n*=19) were published previously [31].

### Statistical analysis

We used the prevalence package in R to estimate global as well as site- and year-specific infection prevalence and 95% confidence intervals (CIs). Next, to identify risk factors for *T. cruzi* infection in vampire bats, we fit three generalized linear mixed models (GLMMs) using the *lme4* package [32]. The first GLMM was fit to our entire dataset (n = 205) and focused only on individual-level predictors, including bat sex, reproductive status, age, and all relevant two-way interactions, given that vampire bats in KK were not sampled in 2021. To next test for site differences in *T. cruzi* infection without confounding site and season, our second GLMM was fit to only data from 2019 and 2022 (n = 191), including site, year, and their interaction as predictors alongside all individual-level main effects and any significant interaction terms from our first model. Lastly, to assess temporal variation in infection status, we fit a third GLMM to just data from LAR (n = 167) with a main effect of year alongside the same individual-level predictors (and interactions) as the previous model. All GLMMs used binomial errors and a logit link, including band number as a random effect, owing to within- or between-year recaptures of 14 individual bats during the study period (representing 30 samples).

### Trypanosoma cruzi *genotyping*

We conducted genotyping on *T. cruzi*–positive samples with sufficient remaining DNA (n = 62) using PCR amplification of the mini-exon intergenic region. Two complementary PCR assays were used to maximize lineage assignment confidence: (i) the classical multiplex mini-exon assay of Souto et al.. [33] and (ii) an additional PCR amplifying a 450–500 bp mini-exon fragment (primers TrypME/TcCH) designed to provide greater phylogenetic resolution and to detect intra-DTU diversity by sequencing [34].

PCR reactions (25 µL) included template DNA, molecular-grade water as a negative control, and reference strains representing known DTUs (WB1 at 0.05 ng/µL and SC43 at 0.01 ng/µL). Amplicons were resolved with 2.0% agarose gels. We selected a subset of 10 high-quality samples for amplicon-based next-generation sequencing (NGS) of the mini-exon locus. This subset provided sufficient read depth for robust haplotype recovery and phylogenetic placement. For each selected sample, amplicons from the three PCR reactions targeting the mini-exon sequence were pooled and purified using the Invitrogen PureLink™ Quick PCR Purification Kit (Cat. No. K3100-02; Thermo Fisher Scientific Baltic UAB, Vilnius, Lithuania) according to the manufacturer’s instructions. DNA concentration of 1 μL of each pooled, purified product was measured with a NanoDrop™ 2000 spectrophotometer (Thermo Scientific™, Wilmington, DE, USA) prior to sequencing.

### Next-Generation sequencing and sequence analysis

Amplicon libraries of the mini-exon gene were prepared following end-repair and indexing, and sequenced on a MiSeq platform (Illumina, Applied Biological Material Inc.) at the Tulane National Primate Research Center. Libraries were multiplexed, and sequencing produced between 134,000 and 650,000 paired end reads per bat blood DNA sample after removal of low-quality and short fragments.

Raw Fastq sequences files were imported into Geneious 11 software for analysis, and quality filtering was applied prior to mapping. Reads were aligned to a curated set of *T. cruzi* mini-exon reference sequences representing each of the seven DTUs (TcI–TcVI and TcBat). Reference sequences used were TcI: H1 (EF576846), TcII: Tu18 (AY367125), TcIII: M5631 (AY367126), TcIV: 92122102r (AY367124), TcV: SC43 (AY367127), TcVI: CL (U57984) and TcBat: TCC2477cl1 (KT305884). To discard the presence of any other trypanosomatides in bats reads were also aligned against the mini-exon sequence of *Trypanosoma dionisii* (AJ250744), a related bat trypanosome within the *T. cruzi* clade. Assemblies to each reference DTU were then analyzed separately. Partial matches to a DTU were discarded from the analysis, and only assemblies covering the full-length of the expected mini-exon PCR products were considered to ensure specificity. Mini-exon sequences were then trimmed of PCR primer sequences [35].Sequence polymorphisms and haplotypes were identified using the FreeBayes SNP/variant tool. Only high-quality alignments were retained, and variants accounting for <1% of total reads per sample were excluded to minimize background noise [36]. Mini-exon sequences from vampire bats have been deposited in GenBank (accessions PX660128-PX660163)

### Phylogenetic analyses

Maximum likelihood phylogenies were built using PhyML as implemented in Geneious [37], and mini-exon sequences from reference parasite strains from all DTUs were included additionally to sequences from triatomine *T. dimidiata* TcI (MW861785), TcIV (MW861768) [17], and human case from Belize TcII (OM818333–34), TcIV (OM818340), TcV (OM818332–41) [18]. Resulting trees were then edited and visualized using FigTree (Interactive Tree of Life) [38].

## Results

### *Prevalence of* Trypanosoma cruzi *in vampire bats*

Across 205 vampire bats sampled in our two sites and over three years, 85 bats (41.5%, 95% CI: 34.9–48.3%) tested positive for *T. cruzi*. The kDNA assay identified more positives (83/205, 40.5%) than the satellite DNA assay (52/205, 25.4%), with concordant amplification in 50 samples. Thirty-three samples were positive only by the kDNA assay, while only two were unique to the satellite DNA assay, consistent with the greater sensitivity of the kDNA target in low-parasitemia samples.

Our first GLMM of individual-level risk factors did not identify effects of age, sex, reproductive status, or their interactions on the probability of *T. cruzi* infection (Table S1, *R^2^* = 0.03, *R^2^* = 0.04). When analyzing only data from 2019 and 2022 to assess spatial effects, we found significant main effects of site and year but not their interaction (Table S2, *R^2^* = 0.16, *R^2^* = 0.18). After adjusting for individual-level covariates, vampire bats had higher odds of *T. cruzi* infection in the LAR (OR = 2.89, *p* = 0.03) and in 2022 (OR = 3.75, *p* < 0.01). When assessing annual variation more explicitly in the LAR in our third GLMM (Table S3, *R^2^* = 0.12, *R^2^* = 0.17), we found that the odds of infection were greater in 2022 than in 2019 (OR = 3.91, *p* < 0.01) and in 2021 (OR = 6.78, *p* = 0.01), with no difference between 2019 and 2021 (OR = 0.58, *p* = 0.41), suggesting annual and seasonal variation in *T. cruzi* infection risks (Figure 2). More generally, these results support widespread infection of vampire bats in northern Belize with *T. cruzi*.

**Figure 2.**
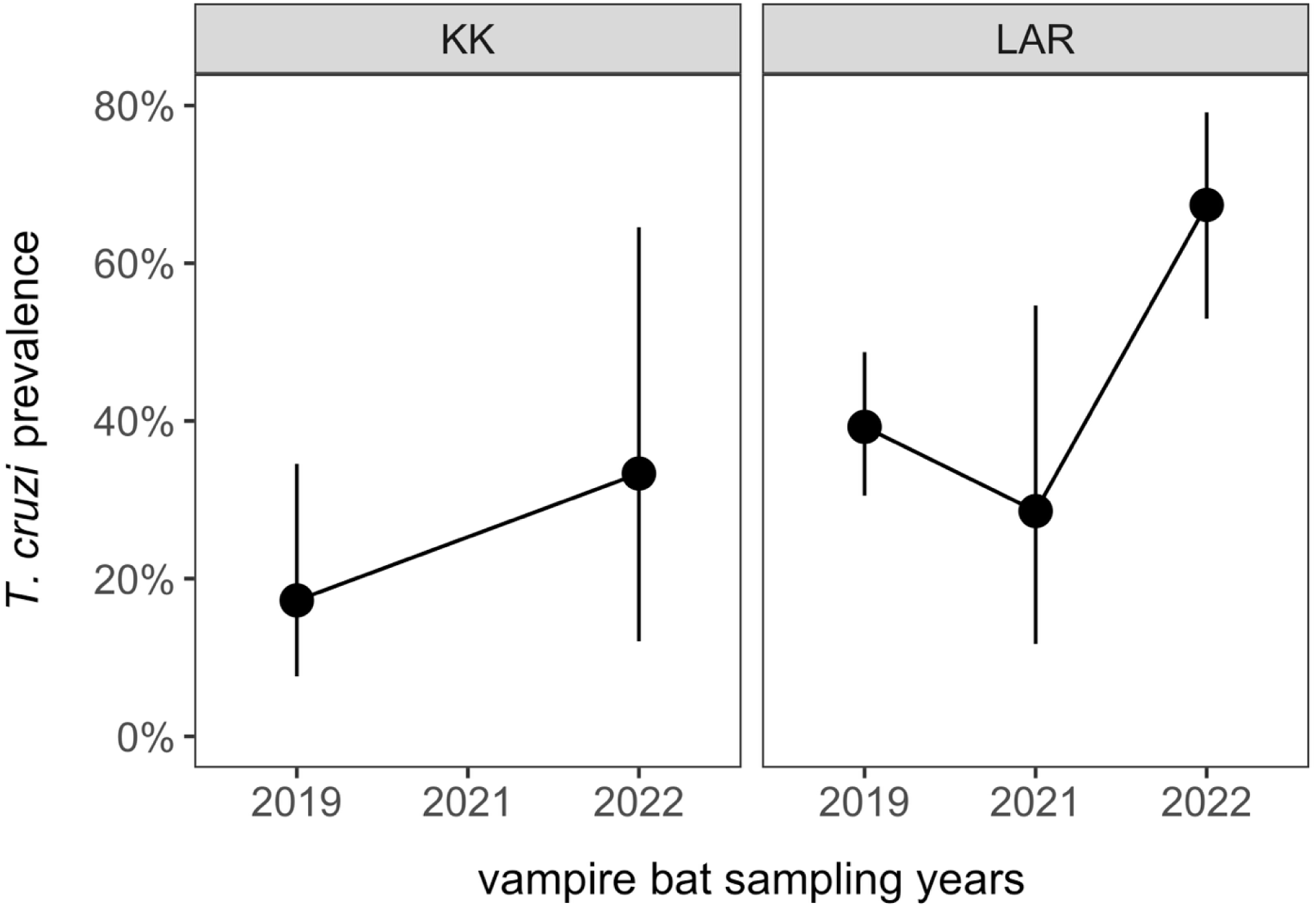
Temporal and spatial variation in *Trypanosoma cruzi* infection prevalence among vampire bats in northern Belize. Point estimates and 95% confidence intervals of *T. cruzi* infection prevalence are shown for (KK) and the LAR across three years (2019, 2021, and 2022).

Across all three GLMMs, individual bat identity had relatively weak contributions to variance explained (i.e., the difference between *R^2^* and *R^2^* ranged from 1–5%). In our small sample of 14 recaptured individuals, eight did not change infection status within or between capture years. Two bats exhibited losses of infection during short timescales (i.e., our two-week sampling periods), three bats gained infection between the wet and dry season (2021–2022), and one bat gained infection between 2019 and 2022 (Figure 3).

**Figure 3.**
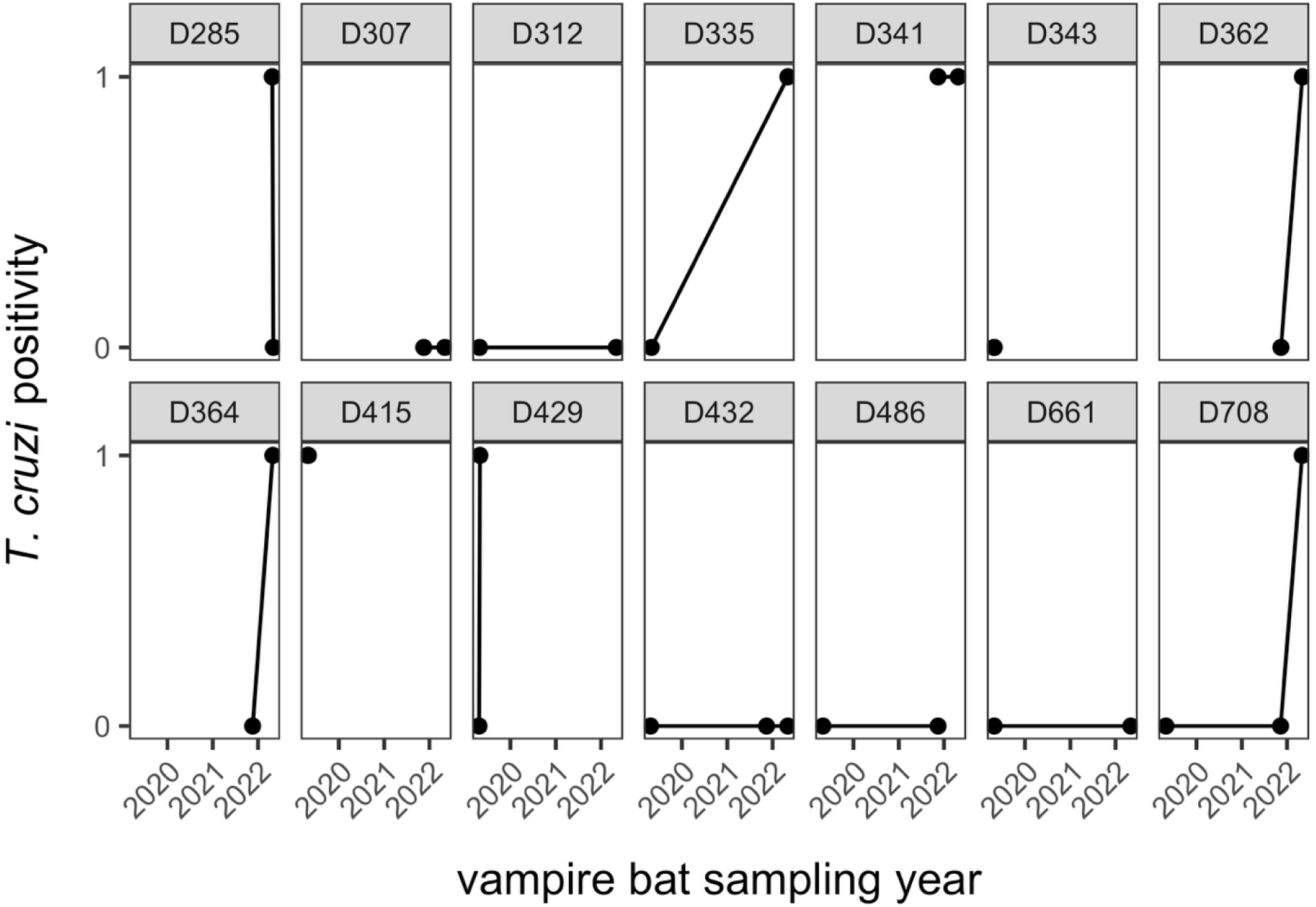
Infection dynamics among resampled vampire bats in northern Belize.

### T. cruzi *sequence diversity in vampire bats*

To refine DTU assignments and characterize intra-lineage diversity, we selected 10 representative *T. cruzi*–positive samples for amplicon-based NGS of the mini-exon locus. Selection criteria included strong PCR amplification, high DNA quality, and representation across multiplex genotyping profiles. For each sample, amplicons from three independent PCR reactions were pooled to maximize haplotype recovery.

NGS analysis showed 36 high-confidence haplotypes across the 10 samples, with bats carrying between 1 and 9 haplotypes per individual, but all haplotypes within a given individual consistently belonged to the same DTU, indicating intra-DTU haplotypic diversity rather than mixed-DTU infections. Based on the maximum-likelihood phylogeny (Figure 4), haplotypes belonged to five DTUs: TcI, TcIV, TcV, TcVI, and TcBat, with TcBat as the predominant lineage. A single divergent sequence grouped robustly within the TcIV clade. No haplotypes corresponding to TcII or TcIII were detected.

**Figure 4.**
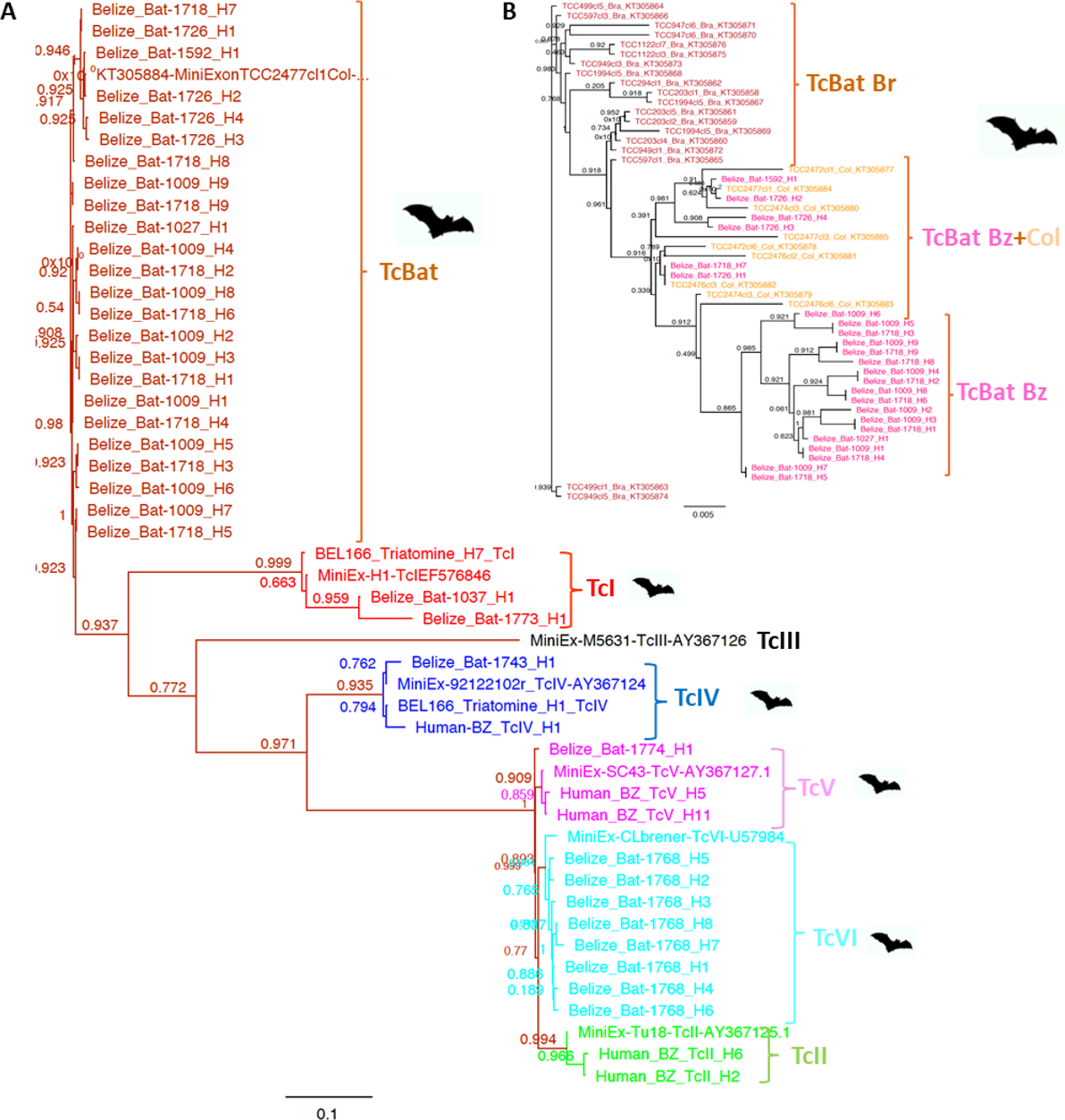
Phylogenetic analysis of *Trypanosoma cruzi* haplotypes recovered from Belizean vampire bats. (A) Maximum-likelihood phylogeny showing that Belizean haplotypes cluster within five DTUs (TcI, TcIV, TcV, TcVI, and TcBat). (B) TcBat haplotypes from Belize form two major groups: one clustering closely with Colombian (Col) TcBat reference sequences, and another forming a distinct Belize-specific clade (Bz). The Brazilian (Br) TcBat lineage is more distantly related. DTUs are color-coded, and bat icons indicate haplotypes generated in this study.

Although individual bats frequently carried multiple haplotypes, all haplotypes within a given bat belonged to the same DTU, demonstrating intra-lineage (rather than mixed-DTU) infections. TcBat included several Belize-specific haplotypes not closely matching available references, representing the first detection of this lineage in Central America. No sequences from other trypanosomatids were recovered.

### *Phylogenetic analysis of* T*. cruzi haplotypes*

Maximum-likelihood phylogenetic analysis of the 36 haplotypes (Figure 4) confirmed their grouping within five DTUs: TcBat (68.6%), TcVI (22.8%), TcI (5.7%), TcIV (2.9%), and a single TcV haplotype. All DTU assignments were supported by clustering with their respective reference sequences.

The TcBat haplotypes identified here formed a distinct, well-supported cluster that branched closest to TcI reference sequences, consistent with the proposed evolutionary origin of TcBat [6, 9, 39, 40]. Within this cluster, several Belizean haplotypes grouped tightly with Colombian TcBat sequences [12], while the Brazilian TcBat lineage was more distantly related. A subset of Belizean haplotypes formed a separate, Belize-specific cluster (Figure 4B).

The single TcIV haplotype identified here clustered with 93% bootstrap support alongside TcIV sequences from *Triatoma dimidiata* collected in Belize and with a strain reported from an acute human infection in 2020, reinforcing the link between sylvatic bat cycles and potential human transmission [17–19].

The eight TcVI haplotypes clustered within the TcVI clade and grouped with the CL Brener reference strain included in the analysis. Although the available reference dataset is limited, the Belizean TcVI haplotypes showed internal diversity within the clade, suggesting intra-DTU variation but not forming distinct sub-lineages (Figure 4A). This observation is similar to a recent report of TcVI in non-human primates [41, 42] and rodents in North America [43, 44],suggesting that this lineage may be more broadly distributed in sylvatic reservoirs than previously appreciated.

Together, these results demonstrate that Belizean vampire bats harbor multiple *T. cruzi* DTUs with substantial haplotypic diversity, including lineages of recognized epidemiological importance.

## Discussion

Our findings provide new insights into the eco-epidemiology of *Trypanosoma cruzi* infections in *Desmodus rotundus* in Central America and northern Belize specifically. With an overall prevalence of 41.5%, common vampire bats are important sylvatic hosts within local *T. cruzi* transmission networks. This overall infection prevalence is consistent with, and in some cases exceeds, those reported in vampire bat populations from Peru, Brazil, and Mexico, where prevalence has been estimated to range from 10–40% [13, 45–48]. These results support growing evidence that vampire bats play a significant role in maintaining *T. cruzi* circulation in wildlife communities and may contribute to sylvatic–domestic spillover dynamics, particularly in areas where human–wildlife interfaces are expanding [11, 47, 49, 50].

In Central America, the contribution of bats to *T. cruzi* transmission has been underexplored, with most research focused on terrestrial mammals (e.g., opossums, armadillos, and rodents) and synanthropic vectors such as *Triatoma dimidiata* [51]. Our study fills a geographic and ecological gap by documenting both high prevalence and substantial parasite diversity in *D. rotundus* from northern Belize, where Chagas disease remains endemic but underdiagnosed [11]. Given that *Desmodus rotundus* frequently feeds on domestic animals and occasionally humans [13, 15], its potential to serve as a host (and possibly a reservoir) as well as a mechanical bridge between sylvatic and domestic cycles warrants further attention [12, 50, 52–54].

A striking result of our study is the high diversity of *T. cruzi* DTUs detected in vampire bats. Haplotypes corresponding to TcI, TcIV, TcV, TcVI, and TcBat were identified using NGS of the mini-exon locus. This level of haplotypes diversity in a single host species, and within a relatively small geographic area, is remarkable and suggests complex transmission networks involving multiple parasite DTUs and vector species. TcI, the most widespread DTU in the Americas, is commonly found in both sylvatic and domestic transmission cycles, including in human cases across Central America [6].

In addition to TcI, TcIV, TcVI, and TcBat, we also detected a TcV haplotype, a lineage typically associated with domestic transmission and human disease in South America [8]. The detection of TcV haplotypes in Belize aligns with recent evidence of a multiclonal human infection involving TcV [18]. Although sylvatic hosts of TcV are considered rare [8, 44], our findings suggest that vampire bats may contribute to the local maintenance of TcV. Identifying the distribution and frequency of TcV in bats and other host taxa will be essential to assess its role in regional spillover or potential spillback from domestic or human transmission cycles given recent evidence of TcV involvement in an acute human case in northern Belize [18]. Resolving the frequency, host range, and directionality of TcV transmission will be critical for understanding its regional significance [17, 18].

TcBat was the most prevalent DTU in our sequenced samples and included several Belize-specific haplotypes, expanding its known geographic range. Belizean TcBat sequences clustered closely with Colombian TcBat references, whereas the Brazilian TcBat lineage was more distantly related, consistent with structured geographic variation. TcBat was long thought to occur predominantly in South American chiropterans [12, 54]. While TcBat has not yet been linked to human disease, its high prevalence raises important questions about its evolutionary origin, host specificity, and ecological function [10, 13]. Likewise, the sympatric presence of TcI, TcIV, TcV, TcVI, and TcBat across the study sites indicates that multiple DTUs circulate within the same bat populations. Although individual bats typically carried multiple haplotypes belonging to a single DTU rather than mixed-DTU infections, the co-circulation of these lineages at the population level could create ecological opportunities for inter-lineage contact in vectors or other hosts. Despite detecting multiple DTUs across the two sampled vampire bat populations here, we found no evidence of mixed DTU infections in individual bats. This contrasts with reports of frequent coinfections in other mammalian hosts and vectors [8, 44, 55].

Taken together, the NGS data indicate that infections in Belizean vampire bats consisted of multiple haplotypes per individual but consistently representing a single DTU, suggesting substantial intra-DTU variation rather than multi-DTU coinfections We also observed spatial and temporal variation in *T. cruzi* infection risk, with higher prevalence at the LAR compared to KK and higher prevalence in 2022 than in 2019 and 2021 (Figure 2). These differences may reflect habitat characteristics, livestock availability, and climate variability, which are known to influence vector populations, bat immunity, and host exposure in vector-borne disease systems [23, 27, 56]. Such patterns may also reflect the impacts of increasing agricultural land conversion on host–host and host–vector contact [57] as well as habitat and seasonal effects on bat immunity [58, 59]. Although our study was not designed to robustly assess individual infection trajectories, we also observed cases of both infection gain and infection loss over relatively short periods; future longitudinal work will be essential to clarify infection persistence and reinfection dynamics of *T. cruzi* in vampire bats (Figure 3).

Our study has some limitations. Sampling was restricted to two sites over three years, only partly assessing seasonal variation, with one roost being abandoned midway through our sampling. Further, only a subset of PCR-positive samples was sequenced, limiting our ability to fully characterize *T. cruzi* diversity. We also did not measure vector abundance, blood meal sources, or host immune function, all of which would provide further insight into the ecological drivers of parasite transmission. Nevertheless, the high prevalence and DTU diversity detected in vampire bats emphasize the complexity of local *T. cruzi* ecology, likely involving diverse hosts, and overlapping transmission pathways. Future studies integrating entomological surveys, blood meal analysis, and parasite genotyping from vectors and other mammals will be essential to construct a more holistic picture of transmission networks in Belize.

From a public health perspective, the detection of TcV and TcVI, DTUs strongly associated with human disease in South America, along with the frequent circulation of TcI, highlights the potential for sylvatic spillovers into domestic environments. Belize remains one of the least studied countries in Central America with respect to Chagas disease, yet recent events indicate that spillover is already occurring: in 2020, an acute case in a child in northern Belize was linked to multiple DTUs (TcII, TcIV, TcV) and to infected triatomines within the household [18] Subsequent entomological work identified a novel *Triatoma* species carrying TcIV from the same household [19]. Moreover, our previous work on *T. dimidiata* in southern Belize revealed clear evidence of local parasite differentiation and circulation of TcI and TcIV in vectors, together with frequent blood-feeding on humans and other mammals, highlighting a highly connected sylvatic–domestic interface in the region [17]. The overlap between DTUs detected in bats, vectors, and humans strongly suggests that vampire bats are part of broader regional transmission networks. Like multiclonal infections detected in *T. dimidiata* in Mexico [55], the diversity found in Belizean vampire bats may represent an overlooked source of epidemiologically relevant strains.

The high prevalence of *T. cruzi in* Belizean vampire bats, together with the observation that individual bats carried multiple haplotypes within single DTUs, suggests repeated exposure to circulating parasite populations. Because *T. dimidiata* rarely feeds on vampire bats and primarily feeds on humans and other mammals in Belize, infection in *D. rotundus* may occur predominantly through feeding on infected prey rather than via vectorial transmission. Comparative data on infection pathways in vampire versus non-vampire bats remain limited, underscoring the need for studies explicitly addressing transmission routes in chiropteran hosts

## Conclusions

Our study demonstrates that common vampire bats in northern Belize harbor a high prevalence of *T. cruzi* and an unexpected diversity of parasite DTUs, including TcI, TcIV, TcV, TcVI, and TcBat. The detection of multiple DTUs in a single host species and geographic area highlights the complexity of transmission networks in this region and underscores that Belize harbors a more dynamic *T. cruzi* landscape than previously recognized.

Similar findings of diverse DTUs in non-human primates, rodents, and other wildlife in North America further emphasize that parasite diversity across the Americas is broader than historically appreciated. These results reinforce the importance of sustained vigilance, since underestimating genetic variability may obscure risks of spillover and transmission to humans.

Given that *Desmodus rotundus* feeds regularly on livestock and occasionally on humans, these bats represent potential bridge hosts that may facilitate spillover between sylvatic, domestic, and human cycles. From a One Health perspective, integrated surveillance that simultaneously monitors bats, vectors, domestic animals, and humans will be essential to anticipate and mitigate the public health consequences of Chagas disease in Central America. In addition, the presence of TcV, a DTU typically linked to domestic transmission and human disease, raises the possibility of spillback from domestic or peridomestic sources into sylvatic bat populations, further underscoring the need for coordinated cross-sector surveillance.

## Acknowledgements

For assistance with bat sampling logistics and research permits, we thank Mark Howells, Neil Duncan, and the staff of the Lamanai Field Research Center.

---

**Supplemental Table 1.** GLMM results for model 1 (individual-level predictors) of *T. cruzi* infection (*n* = 205).

| Predictor | $\chi^2$ | df | $p$ |
| --- | --- | --- | --- |
| Age | 0.01 | 1 | 0.91 |
| Sex | 1.55 | 1 | 0.21 |
| Reproductive status | 0.62 | 1 | 0.43 |
| Age * sex | 1.89 | 1 | 0.17 |
| Sex * reproductive status | 0.41 | 1 | 0.52 |

**Supplemental Table 2.** GLMM results for model 2 (spatiotemporal predictors) of *T. cruzi* infection (*n* = 191).

| Predictor | $\chi^2$ | df | $p$ |
| --- | --- | --- | --- |
| Age | 0.01 | 1 | 0.93 |
| Sex | 0.74 | 1 | 0.39 |
| Reproductive status | 0.28 | 1 | 0.60 |
| Site | 4.76 | 1 | 0.03 |
| Year | 10.08 | 1 | 0.002 |
| Site * year | 0.06 | 1 | 0.80 |

**Supplemental Table 3.** GLMM results for model 3 (temporal predictors) of *T. cruzi* infection (*n* = 167).

| Predictor | $\chi^2$ | df | $p$ |
| --- | --- | --- | --- |
| Age | 0.20 | 1 | 0.65 |
| Sex | 0.90 | 1 | 0.34 |
| Reproductive status | 1.45 | 1 | 0.23 |
| Year | 10.12 | 1 | 0.006 |

